# smORFer: a modular algorithm to detect small ORFs in prokaryotes

**DOI:** 10.1101/2020.05.21.109181

**Authors:** Alexander Bartholomäus, Baban Kolte, Ayten Mustafayeva, Ingrid Goebel, Stephan Fuchs, Susanne Engelmann, Zoya Ignatova

## Abstract

Emerging evidence places small proteins (≤ 50 amino acids) more centrally in physiological processes. Yet, the identification of functional small proteins and the systematic genome annotation of their cognate small open reading frames (smORFs) remains challenging both experimentally and computationally. Ribosome profiling or Ribo-Seq (that is a deep sequencing of ribosome-protected fragments) enables detecting of actively translated open-reading frames (ORFs) and empirical annotation of coding sequences (CDSs) using the in-register translation pattern that is characteristic for genuinely translating ribosomes. Multiple identifiers of ORFs that use 3-nt periodicity in Ribo-Seq data sets have been successful in eukaryotic smORF annotation. Yet, they have difficulties evaluating prokaryotic genomes due to the unique architecture of prokaryotic genomes (e.g. polycistronic messages, overlapping ORFs, leaderless translation, non-canonical initiation etc.). Here, we present our new algorithm, smORFer, which performs with high accuracy in prokaryotic organisms in detecting smORFs. The unique feature of smORFer is that it uses integrated approach and considers structural features of the genetic sequence along with in-register translation and uses Fourier transform to convert these parameters into a measurable score to faithfully select smORFs. The algorithm is executed in a modular way and dependent on the data available for a particular organism allows using different modules for smORF search.

## INTRODUCTION

Next-generation sequencing (NGS) technologies enable a rapid and easy detection of genomic information of new species. However, delineating the protein-coding open reading frames (ORFs) in genomes after sequencing and *de novo* genome assembly remains still a challenge. After the pioneering effort of Fickett to unify concepts on how to define protein-coding sequences (1), further criteria have been added to increase the confidence in *de novo* identifications. These include intrinsic signals involved in gene specifications (e.g., start and stop codon, splice sites), conservation patterns in related genomes with weighted conservation depending on evolutionary distance and verification to known ORFs or protein sequences (2,3). Classically, these rules in the genome annotation protocols are performing well only on larger ORFs which span at least 100 codons (4,5), thus small ORFs (smORFs) shorter than 100 codons are systematically underrepresented and cannot be identified by common algorithms (6). Mounting evidence suggests crucial functions for smORFs in cellular and molecular processes in both eukaryotes (6-13) and prokaryotes (14-22). However, systematic identification of functional small proteins or microproteins (also called micropeptides) remains challenging both experimentally and computationally.

Recent developments of the NGS technologies to probe the position of translating ribosomes with codon precision – ribosome profiling or Ribo-Seq (23), enable detecting actively translated ORFs by capturing ribosome-protected fragments (RPFs) and empirically annotate coding sequences (CDSs). Several new previously unannotated ORFs, including smORFs, have been identified mostly in eukaryotes (8,24-26). Some studies oppose that RPFs alone are not sufficient to classify a transcript as protein-coding or non-coding (27). Alternatively, Poly-Ribo-Seq which specifically sequences polyribosomes separated through sucrose gradients is suggested as more stringent approach in isolating translated ORFs (28). mRNAs translated by more than one ribosome (i.e. polyribosomes) are classically defined as genuinely translated mRNAs. However, given that a ribosome protects on average of 26-30 nt this approach may miss very short smORFs (less than 10 amino acids) whose size might permit translation by a single ribosome and thus migrate in the monosomal fraction. Ribo-Seq combined with a treatment with antibiotic that specifically stalls ribosomes at translation initiation site (TIS-Ribo-Seq) selects for potential new initiation sites and allows detecting new ORFs in non-coding regions or overlapping ORFs which overlap with annotated ORFs and are undistinguishable in the Ribo-Seq data sets (8,19,22,25,29-32).

Complementing Ribo-seq with computational predictions revealed several hundred smORFs in eukaryotes (8,24,26,33,34). The crucial metrics they use is the enrichment of RPFs in the ORFs and the 3-nt periodicity characteristic for genuinely translating ribosomes. These approaches have difficulties evaluating prokaryotic genomes due to their unique architecture, including polycistronic messages, large fraction of overlapping ORFs, leaderless translation and lack of classical ribosome-binding site (i.e. with direct start of translation from the start codon (35,36)). The resolution of the prokaryotic Ribo-Seq sequencing data is lower than in eukaryotes due to intrinsic properties of the nucleases used to generate the RPFs (37), which often results in imperfect periodicity. Together, this makes a genome-wide identification of smORFs encoding functional small proteins in prokaryotes even more challenging.

Here, we present a new algorithm, smORFer, for identifying smORFs by integrating genomic information, structural features, Ribo-Seq and TIS-Ribo-Seq to faithfully select translated and initiated ORFs, respectively. The algorithm is executed in a modular fashion and the use of various modules can be selected dependent on the data availability for each organism. smORFer is versatile and suitable for every organism, but shows high confidence of predictions in particularly difficult-to-annotate smORFs in bacteria.

## MATERIALS AND METHODS

### Data sets used in the analysis

We generated two biological Ribo-Seq replicates for *Staphylococcus aureus* and downloaded *Escherichia coli* MG1655 (Ribo-Seq, GSM3455899 and retapamulin-treated TIS-Ribo-Seq, GSM3455900 (19)) and *Bacillus subtilis* data (Ribo-Seq, GSM872395 and GSM872397, (38)) from the Gene Expression Omnibus (GEO) repository. The Ribo-Seq data for *S. aureus* were uploaded in GEO under accession number GSE150601.

### Ribo-Seq of *S. aureus*

Cells grown in complex medium to OD_550_=1 were harvested by rapid centrifugation, resuspended in ice-cold 20 mM Tris lysis buffer pH 8.0, containing 10 mM MgCl_2_ x 6 H_2_O, 100 mM NH_4_Cl, 0.4 % Triton-X-100, 4 U DNase, 0.4 µl Superase-In (Ambion), 1 mM chloramphenicol and disrupted by homogenisation (FastPrep-24 ™, MP Biomedicals) with 0.5 ml glass beads (diameter 0.1 mm). 100 A_260_ units of ribosome-bound mRNA fraction were subjected to nucleolytic digestion with 10 units/μl micrococcal nuclease (Thermofisher) in buffer with pH 9.2 (10 mM Tris pH 11 containing 50 mM NH_4_Cl, 10 mM MgCl_2_, 0.2% triton X-100, 100 μg/ml chloramphenicol and 20 mM CaCl_2_). The rRNA fragments were depleted using the *S. aureus* riboPOOL rRNA oligo set (siTOOLs, Germany) and the library preparation was performed as previously described (39).

### Data processing and mapping

Raw sequencing reads were trimmed using FASTX Toolkit (quality threshold: 20) and adapters were cut using cutadapt (minimal overlap of 1 nt). The following genome version were used for mapping: *E. coli* U00096.3, *S. aureus* NC_009641.1 and *B. subtilis* NC_000964.3. Genomes and annotations were downloaded from NCBI (January 2020). In the first step of mapping, reads mapping to rRNAs were discarded. Thereafter, reads were uniquely mapped to the reference genomes using Bowtie (40), parameter settings: -l 16 -n 1 -e 50 -m 1 --strata –best y. Non-uniquely mapped reads were discarded. The total number of mapped reads are summarized in Table S1.

### Workflow

The workflow of smORFer, which is executed in a modular way, is summarized in Fig. 1. Several simple filtering steps are performed using BEDTools (41). Module A is required to define the boarders of all putative ORFs and is refined by the structural properties that are intrinsic to protein coding sequences. Modules B and C add more confidence to the detected smORF candidates and can be executed either independently or together; the latter increases the detection of true positive novel smORFs.

**Figure 1.**
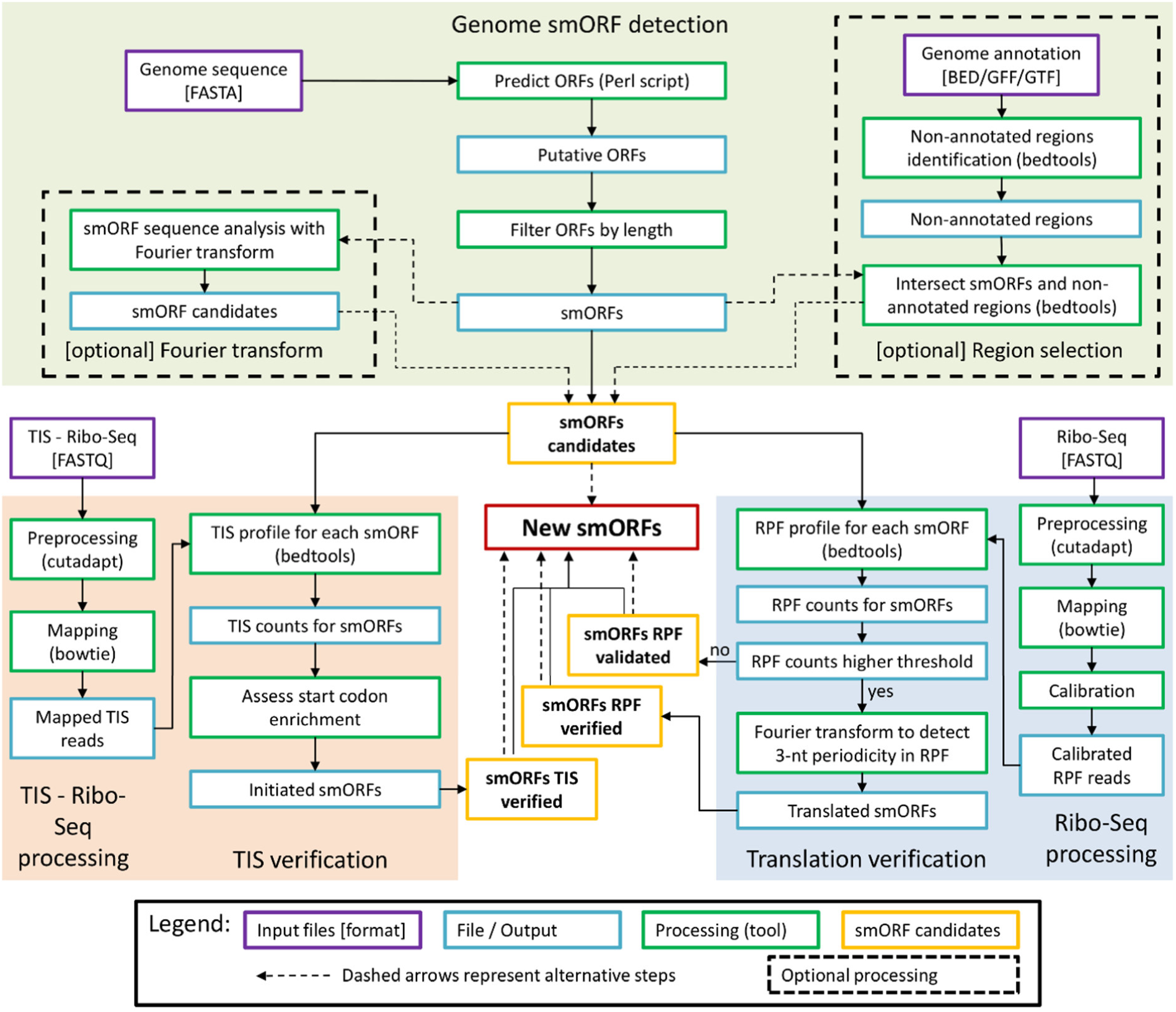
General scheme of smORFer algorithm with its three modules that evaluate genomic information (module A, green), translation or enrichment of RPFs and 3-nt periodicity from Ribo-Seq data (module B, blue), and TIS from TIS-Ribo-Seq data to identify TIS (module C, orange).

#### Genome-based ORF detection (Module A)

The list of putative ORFs was generated using modified Perl script (42); it generates putative ORFs with in-frame start and stop codon. We used four start codons, ATG, GTG, TTG, CTG, which are the most common in prokaryotes (43) and the three uniform stop codons, TGA, TAG, TAA. smORFer separated smORFs based on their location, in the non-annotated and annotated regions, and also contains strand-specific filter for selecting this region.

To detect putative smORFs potentially coding peptides or proteins, i.e. exhibiting 3-nt structural periodicity in the CDS and thus be potentially translated, we use Fourier transform (FT; implemented as R’s base fft function) of the GC content on a single gene basis, i.e. for each single ORF this is a vector of 0’s and 1’s. The signal is first normalized by the ORF length as the signal intensity depends on the ORF length. In the 3-nt periodic pattern the periods of 3 and 1.5 are always present regardless of the length of the putative ORF. Thereafter, we build the fraction of normalized signal at the period of 3 divided by the arithmetic mean of the signal between both periods 3 and 1.5. ORFs with FT≥3 are defined as potentially translated.

#### Detection of translated ORFs from Ribo-Seq data including read processing (Module B)

Ribo-Seq data are mapped and the RPFs are calibrated as described (44); all scripts are available here: https://github.com/AlexanderBartholomaeus/MiMB_ribosome_profiling. smORFs with minimum 5 RPFs are selected and assigned as *validated*. The reliable minimum read counts per gene should be determined individually for each Ribo-Seq using variability analysis of the counting statistics of two independent biological replicates that also assesses the influence of counting noise (23,39). RPF counts ≥ 5 is on average above the counting error for short ORFs in Ribo-Seq data sets and can be used as an arbitrarily set minimum when biological replicates are not available.

The calibration assigns in each read the codon at the ribosomal A or P site correctly, allowing tracking the codon-wise periodic movement of the ribosomes on an ORF. smORFs with a 3-nt periodicity are classified as verified. To position a read in the A or P site of the translating ribosome, the reads are separated by length and the offset is determined for each length (44). Determining the offset for each RPF length individually significantly improves the resolution and the 3-nt periodicity in prokaryotic Ribo-Seq data sets (44) The calibration requires a good read coverage, hence smORFs with a coverage of 100 RPFs per kilobase of ORF length (RPK) we subject to FT analysis to determine the 3-nt periodicity in the calibrated RPF profile. Usually a coverage of 100 RPK (i.e. 1 read per 10 nt) results in good FT analysis.

Similar to the FT analysis of the structural periodicity of each ORF (see above), here the 3-nt periodicity of the calibrated RPF profile is compared to the mean of the signal between the periods of 3 nt and 1.5 nt. smORFs with a value higher of 2 classifies them as verified with genuine translation. smORFs not showing a 3-nt periodic signal, i.e. low RPF coverage, are sorted as validated. Note, that the validated smORFs should be also kept as they are likely to be true hit; they might be translated at low level, which precludes designation of the few reads to a real 3-nt periodic pattern and the FT analysis.

#### Detection of TIS (Module C)

Ribo-Seq is performed in the presence of translation initiation inhibitor, here retapamulin for the *E. coli* data set (19). TIS-Ribo-Seq was preprocessed the same way as Ribo-Seq. From the processed TIS-Ribo-Seq RPFs the middle nucleotide of each read is extracted and used in further analysis. It should be noted that these reads cannot be calibrated, because of the biased coverage at initiation. For each smORF, the middle-nucleotide counts either over the three nucleotides of the start codon or spanning the initiation codon and one codon upstream and downstream of it are summed up and ORFs with more than 5 counts are classified as having a true TIS.

### Operating system and R versions and packages

We used Ubuntu 18.04 LTS as the operating system. For data analysis and visualization, we used R (3.5.0) including packages seqinr (3.6-1) and Biostrings (2.50.2) which are available on all operating system.

## RESULTS AND DISCUSSION

### Design of the smORFer – a modular algorithm to detect smORFs

The availability of various sequencing data (DNA-Seq, Ribo-Seq, TIS-Ribo-Seq) for different organisms may largely vary, hence we sought to develop an algorithm – smORFer – with a modular design which specifically targets the detection of smORFs. smORFer combines three modules which utilize different inputs and can be used independently or in combination to increase the confidence in smORF annotation (Fig. 1). The three inputs are: (1) for module A ‘Genome-based smORF detection’ the input is the genomic nucleotide sequence, (2) for module B ‘*Detection of translated ORFs’* the input is Ribo-Seq data set; and (3) for module C ‘*Detection of TIS*’ the input is TIS-Ribo-Seq (Fig. 1).

#### A. Genome-based ORF detection

This module uses genomic data as an input to first detect ORFs in length-independent manner. In all three organisms tested, we detected a large number of putative smORFs of three codons or longer but smaller than 50 codons. We restricted the maximal length cutoff to ≥ 50 amino acids or codons, which length is defined for the category of small or micropeptides (22,45); the algorithm can perform calls for ORFs at any length. We used four start codons, ATG, GTG, TTG, CTG, which are the most common in prokaryotes (43) and the three uniform stop codons, TGA, TAG, TAA. A single amino acid, although theoretically possible to be produced from a start/stop-ORF (19) does not fulfil the criteria for a peptide and were not considered. Analysis of the genomes from the three organisms revealed a well-defined 3-nt periodic structure within the CDSs with first nucleotide being the most structured (Fig. 2), which is conserved feature of the CDS only (46). Moreover, structural analyses of the transcriptomes shows a conservation of this periodicity among the protein-coding mRNAs in prokaryotes (39), yeast (47), plants (48) and humans (49). Thus, we included a step to assess the discrete *3-nt structural periodicity* of each putative ORF using Fourier transform (FT>3, Fig. 1).

**Figure 2.**
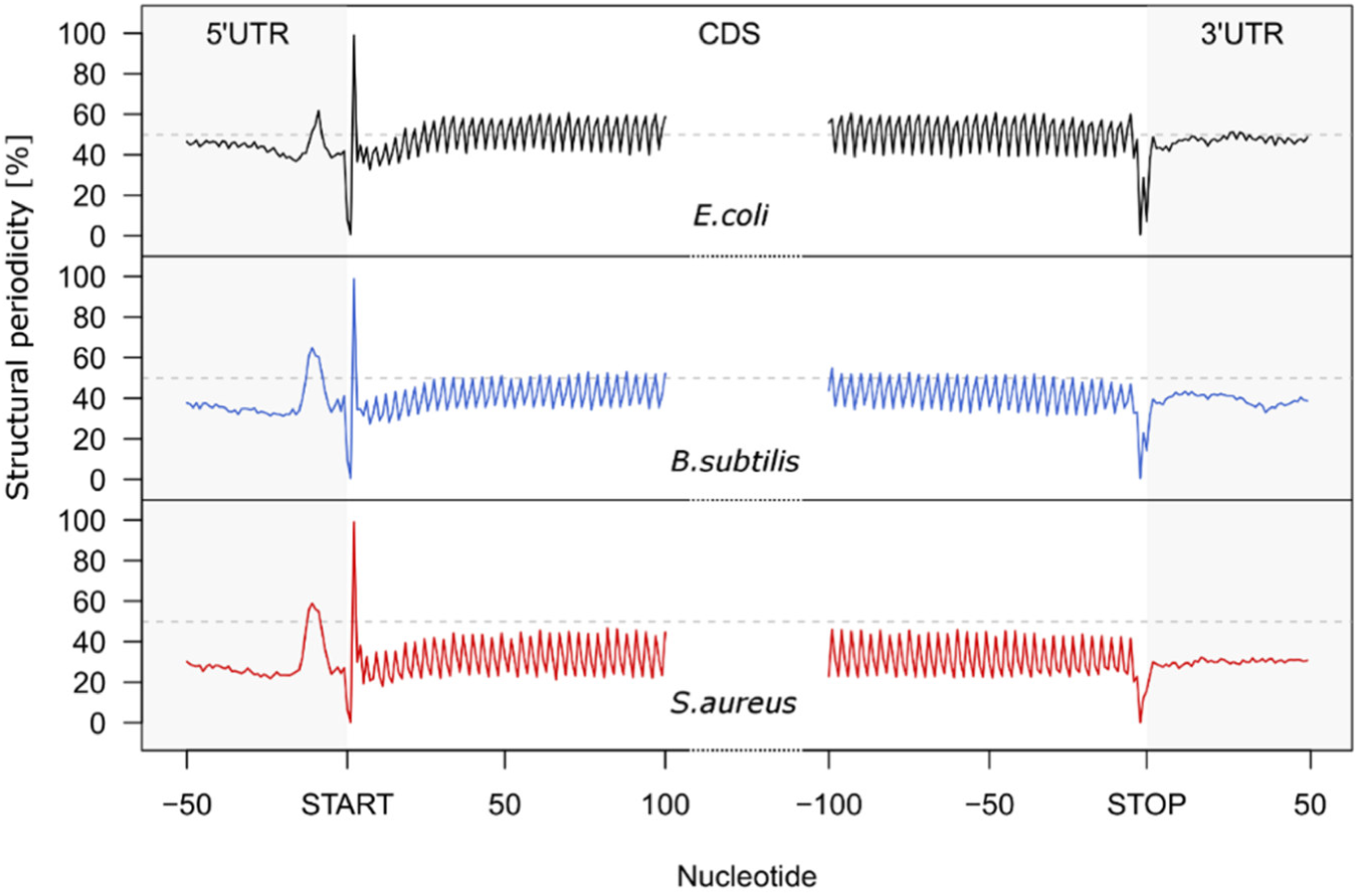
Metagene analysis of the average genomic structure across the 5’UTRs, CDSs and 3’UTRs of all protein-coding transcripts in *E. coli* (A), *B. subtilis* (B) and *S. aureus* (C). ORFs are aligned at the start or stop codon, respectively. Note that the GC content of the species differs and is 51% for *E. coli*, 44 % for *B. subtilis* and 33% for *S. aureus*. Only non-overlapping protein-coding ORFs are plotted. The horizonal dahsed line denotes the average structure of a hypothetical genome with 50 % GC content.

#### B. Detection of translated ORFs from Ribo-Seq data

This module assesses the translation of each ORF from a Ribo-Seq data set (Fig. 1). First, to filter out ORFs with a translation level below a threshold level of sporadic expression, smORFer detects ORFs with a minimal number of RPFs (≥5 RPFs) and categorizes them as *validated*. Genuinely translated ORFs exhibit a 3-nt periodicity in their RPF coverage, hence at a second stage, ORFs undergo a 3-nt periodicity analysis which is assessed again using FT. smORFs over the threshold (FT>2) are categorized as *verified*. Usually the 3-nt pattern is well detectable in smORFs with a good coverage (higher than 100 RPKM, (44)), yet we do not discard the smORFs with no discernible periodic RPF coverage (*validated* category) as they can be still expressed but translated at low level.

#### C. Detection of TIS

This module uses as input TIS-Ribo-Seq data. The following antibiotics have been used so far in prokaryotes retapamulin (19), Onc112 (22), tetracycline (31) to block the transition of the ribosomes from initiation into elongation, thus enabling detection of initiating ribosomes. The smORFer selects positions with minimal number of RPFs covering the start codon (Fig. 1).

### Performance of smORFer for *de novo* identification of smORFs

Here, we employed the smORFer in predicting smORFs in three different organisms, *E. coli, B. subtilis* and *S. aureus*. For all three Ribo-Seq data were available, and TIS-Ribo-Seq only for *E. coli*. In all three organisms tested, based on the genomic sequence and using the first search criterium we detected a large number of putative smORFs of three codons or longer but smaller than 50 codons (>300,000, Table 1). Among the smORFs selected with this very simple selection feature (42), a large portion of them were overlapping, i.e. with different start codons but terminated by one stop codon. Four different start codons, ATG, GTG, TTG and CTG, the most used in bacteria were used as selection criterium. Thereby their usage largely differs by several orders of magnitude, e.g. in *E. coli* the usage is ATG - 81.8%, GTG - 13.8%, TTG – 4.34% and CTG – 0.024% (43). This start codon usage is deduced from the annotated (large) ORFs, but since smORFs may follow non-canonical rules in initiation (35,36), including non-canonical start codons, we kept all those codons with equal weight in the search.

**Table 1.** Comparison of the searches for smORFs in three different organisms for which different data sets were available. For *E. coli* MG1655 Ribo-Seq and TIS-Ribo-Seq are available, whereas for *B. subtilis* and *S. aureus* only Ribo-Seq is available. Module color coding is as in Fig. 1.

| Module | Description of filtering step | *E. coli* | *S. aureus* | *B. subtilis* |
| --- | --- | --- | --- | --- |
| Genome-based smORF detection (module A) | smORFs length $\geq 9$ to $\leq 150$ nt | 415,133 | 305,549 | 405,321 |
|  | smORFs in non-annotated regions | 234,399 | 179,679 | 238,446 |
|  | 3-nt structural periodicity detected by Fourier transform | 10,689 | 9,608 | 13,619 |
| Detection of translated ORFs (module B) | *validated* smORFs with $\geq 5$ RPF | 3,079 | 6,595 | 2,105 |
|  | *verified* smORFs with 3-nt RPF periodicity detected by Fourier transform | 175 | 555 | 168 |
| Detection of TIS (module C) | with $\geq 5$ RPFs at start codon | 424 | n.a. | n.a. |
| Cross comparison | *validated* smORFs and TIS-Ribo-Seq signal | 160 | n.a. | n.a. |
|  | *verified* smORFs with 3-nt RPF periodicity and TIS-Ribo-Seq signal | 16 | n.a. | n.a. |

The coding sequences (CDS) of all three organisms are more structured than the 5’ or 3’UTRs (Fig. 2) and this is independent on the GC content of the organism. Even the most AT-rich genome tested here (*S. aureus*, 33% GC content) exhibits the exactly same structural pattern. Thus, we reasoned that smORFs if protein coding would share the same structural periodicity like the annotated CDSs encoding large proteins. We subjected all smORFs to Fourier transform analysis which converts this characteristic pattern into a score (Fig. 3) and used it for further filtering criterium in module A (Table 1). smORFs because of their short length and higher signal-to-noise ratios (50) are selected with with a FT score higher than 3 to be grouped as potentially translated (Fig. 3C). This parameter significantly reduced the number of the potential candidates, yet, the number of smORFs remained relatively large (Table 1). A validation of this step with annotated ORFs revealed a detection rate of 80%.

**Figure 3.**
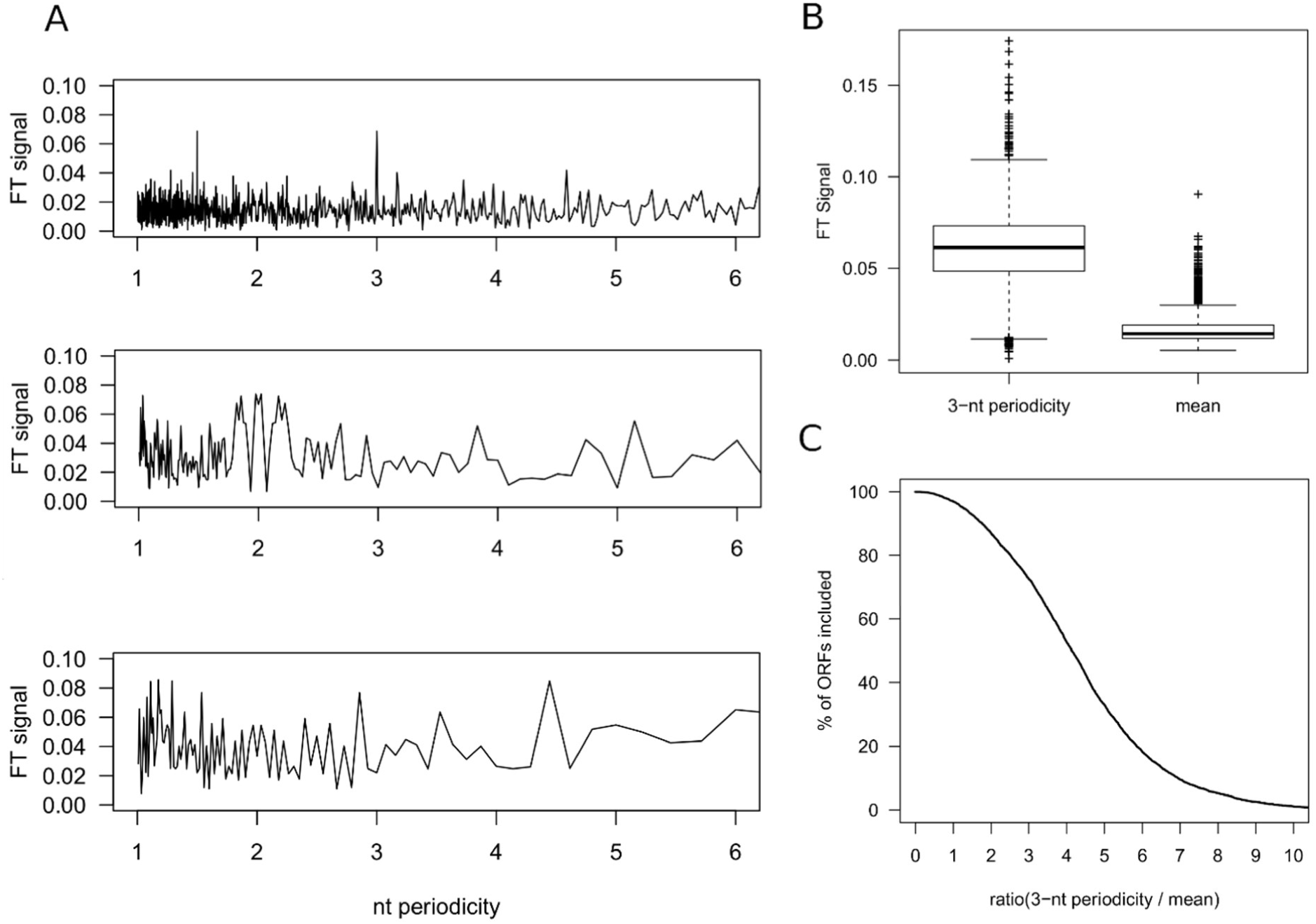
Fourier transform (FT) of the 3-nt structural periodicity in ORFs. (A) Averaged 3-nt structural periodicity of protein-coding ORFs (upper panel), intergenic region (middle panel) and non-protein coding gene (5S rRNA, lower panel). (B) The FT signal normalized by the ORF length at 3-nt periodicity (left) and by the arithmetic mean of the signal between periodicity at 1.5 and 3 (right) for protein coding ORFs. (C) Commulative distributions of averaged FT values (from B) for protein-coding ORFs. It should be noted that a cutoff of 3 detects appr 70% of known ORFs, while a cutoff of 2 – 85% of the known ORFs.

Again, the most of the detected smORFs in this step remained overlapping, i.e. with distinct start codons but terminated by the same stop codon. Within these, the distribution of the smORFs initiated with ATG, GTG, TTG and CTG was for *E. coli* – 900, 625, 815 and 739, respectively, for *B. subtilis* - 702, 422, 609 and 372, respectively, and for *S. aureus* – 2,356, 1,212, 2,227 and 800, respectively. The distribution among the start codons in the putative smORFs is relatively balanced between the four most common codons unlike the highly skewed distribution among them in the long annotated ORFs (43). Yet, at this stage, in order not to miss non-canonically initiated smORFs we do not apply further selection criteria.

Next, using Ribo-Seq data we analyzed the translation status of the smORFs with structural periodicity (module B, Fig. 1). 3,079, 6,595 and 2,105 smORFs for *E. coli, S. aureus* and *B. subtilis*, respectively, were selected with RPFs over the threshold (*validated* candidates, Table 1). Most of which were expressed at very low level with only few RPFs; however, those are still kept for further analysis to not discard lowly expressed smORFs. More stringent criterium for selecting genuinely translated ORFs is assessing their 3-nt periodicity. For this, the RPFs need to be precisely positioned within ORFs, or calibrated by aligning their 5’ or 3’ ends to start or stop codons (51) – a key step to obtain codon resolution and extract 3-nt periodicity, which is a characteristic feature of a genuine translation. All smORFs with sufficient coverage after calibration of the RPFs were subjected to Fourier transform analysis which converts this 3-nt characteristic pattern into a score and smORFs with FT≥2 were defined as *verified* candidates (Table 1). In this step, 175, 555 and 168 non-annotated smORFs were discovered in *E. coli, S. aureus* and *B. subtilis*, respectively (Table 1).

For *E. coli* a TIS-Ribo-Seq data set was available, which we used for further verification of both *validated* and *verified* categories in module B (Table 1). Initially we used a much broader sequence of RPFs coverage to identify TIS, i.e. RPF counts over the threshold on at least one nucleotide in a stretch of 9 nt spanning the start codon and one codon upstream and downstream of it. This search revealed many false-positives (Fig. S1). Inspection of the annotated protein-coding ORFs shows that retapamulin crisply stalls over the start codon, i.e. a true TIS exhibits a maximum centered coverage over the start codon. Therefore, we restricted the calling over the start codon only and from the 3,079 validated smORFs, 160 possessed a TIS signal and from the 175 verified – 16 (Table 1), implying the importance of using various data sets to increase stringency and confidence in smORF identification.

The algorithm successfully detected all experimentally verified smORFs (Fig. 4A) and also some recently identified smORFs with manually assessed TIS (19,22) (Fig. S2). In addition, smORFer also detected some new smORF candidates (Fig. 4B). The TIS-Ribo-Seq allows to precisely position the likely true start codon (Fig. 4B) in the overlapping frames, i.e. such with different initiation codon but common stop codon, which are counted as independent smORFs in the modules A and B (Table 1). It is worth mentioning, that retapamulin is so far the only initiation inhibitor for gram-positive bacteria, which exhibits such precise inhibition (19) and allows for exact detection of TIS. Other antibiotics show much broader coverage across initiation sites and are not always precisely centered at the initiation codon (22,31). To decrease the false-positive hits, in particular for very short smORFs, we still recommend executing restrictive call with a coverage over the start codon and expand it by maximally one nucleotide at each site.

**Figure 4.**
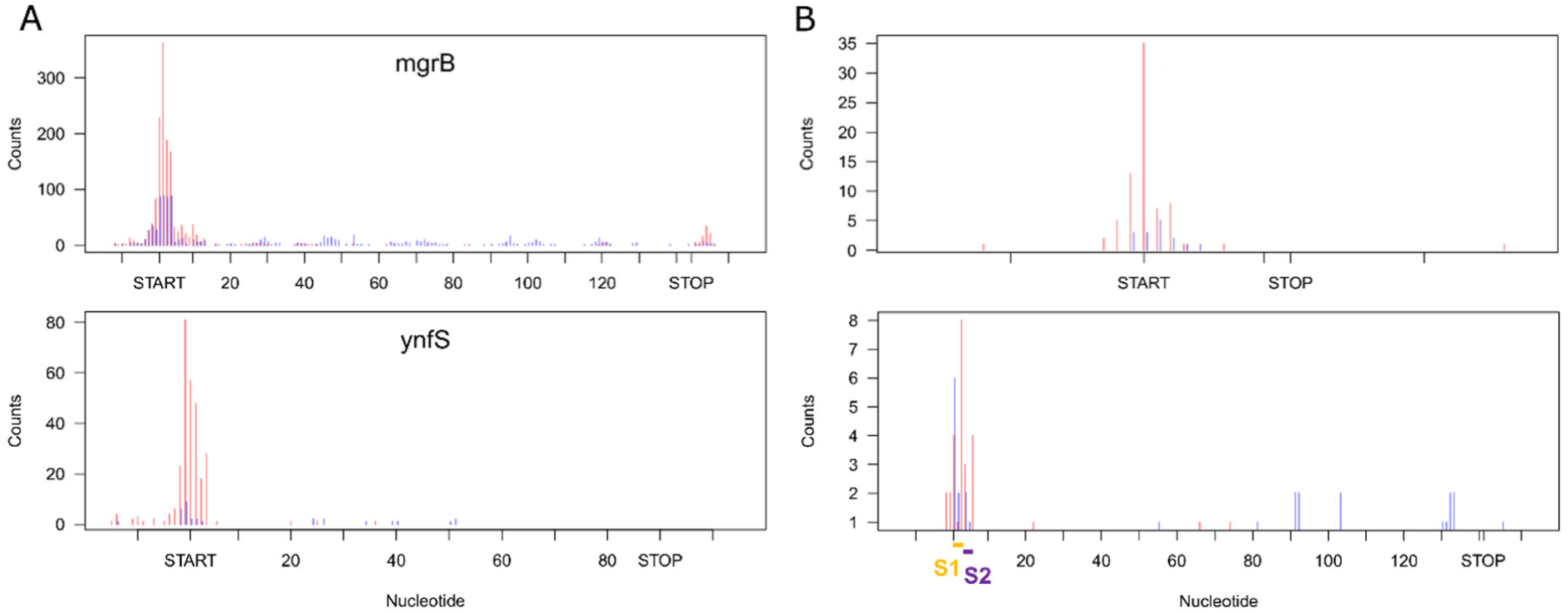
Examples of smORFs detected in *E. coli* with smORFer. (A) All experimentally verified smORFs are detected with smORFer. (B) Examples of two newly identified smORFs with an example for a member of the validated (only RFP coverage) and verified (3-nt periodicity in the RPF coverage) categories. The red counts represent the counts from TIS-Ribo-Seq and blue from the Ribo-Seq. S1 and S2 denote two start codons detected in these two overlapping smORFs sharing the same stop codon, and TIS-Ribo-Seq proves the first one (S1) to be true initiation site. Note that the signal in the TIS-Ribo-Seq with retapamulin as TIS inhibitor is precisely centered at the start codon.

In all three categories, total, verified and validated smORFs, which were verified in the TIS-Ribo-Seq data set for *E. coli* (Table 1), we analyzed the distribution of the start codons. Among the 424 smORFs with TIS signal the distribution of the smORFs initiated with ATG, GTG, TTG and CTG was – 229, 65, 92 and 38, respectively. While for the 160 verified smORFs the distribution of initiation codons was similar – 79, 24, 43 and 14, respectively, for the validated 16 candidates this changed to – 5, 1, 7 and 3, respectively. This different usage of start codons within the most stringent – the validated group, suggests that smORFs exhibit a different bias of start codon usage than long protein-coding ORFs (43). It is conceivable based on this distribution to include a start-codon selection step in module A; yet, data are available for only one single organism and ideally this distribution, if uniform, should be verified for other bacteria.

## Conclusion

Comprehensively designed for annotating *de novo* smORFs using various data sets, smORFer presents remarkable advantages. It has a high efficiency in predicting smORFs with high probability to be expressed. The modularizable structure of smORFer offers advantages in verifying smORFs calling dependent on the available data sets for each organism. The higher the number of the data sets and the modules run in smORFer, the higher the accuracy of the smORF prediction. Many smORFs might be expressed only under stress conditions. Hence the next challenge is to surgically dissect their expression with Ribo-Seq and TIS-Ribo-Seq collected under various stress conditions. This will allow conditionally translated smORFs to be disambiguated from the pool of smORFs with no RPFs under permissive conditions, i.e. categorized as untranslated. When paired to smORFer such data sets, expression events, even conditional expression events, will be mapped more comprehensively.

Computationally, smORFer enables full analysis in a standardized way requiring little computational resources. The workflow in each module is easy to use and simple to modify to achieve high precision in smORFs calling.

## SUPPLEMENTARY FIGURES AND TABLES

**Table S1.** Mapping outcome for different data sets.

| Organism | Total number of sequencing reads | Uniquely mapped reads |
| --- | --- | --- |
| *E. coli* strain MG1655 (19) |  |  |
| Ribo-Seq | 65,014,082 | 65,014,082 |
| TIS-Ribo-Seq | 36,693,777 | 6,534,911 |
| *S. aureus* strain Newman (this work) |  |  |
| R#1 Ribo-Seq | 26,221,544 | 3,366,465 |
| R#2 Ribo-Seq | 20,114,374 | 2,334,161 |
| *B. subtilis* strain 168 (38) |  |  |
| R#1 Ribo-Seq | 18,230,224 | 6,807,640 |
| R#2 Ribo-Seq | 35,407,892 | 12,531,457 |

**Figure S1.**
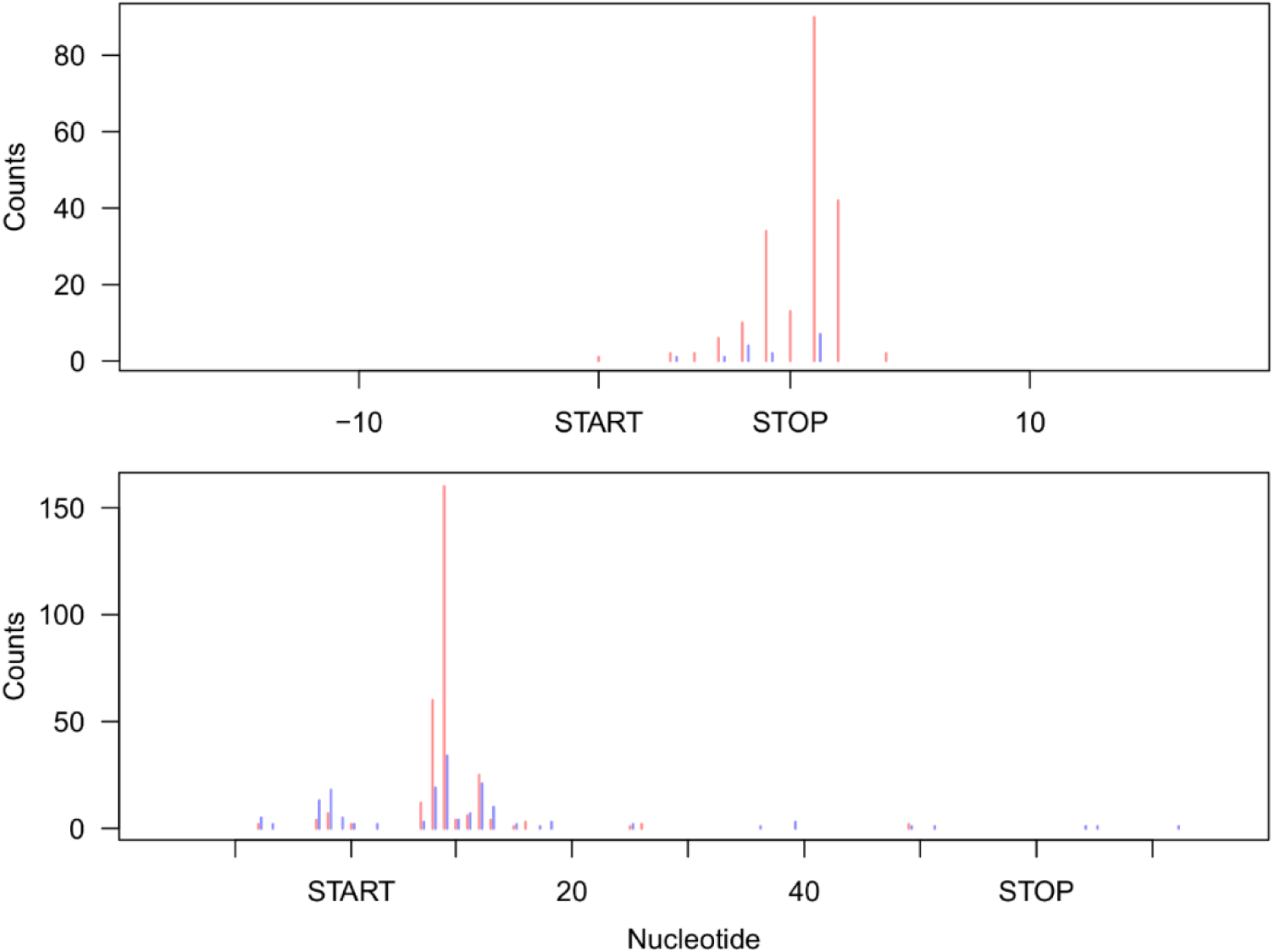
Considering TIS coverage over three codons results in some false-positives. Examples of two newly identified smORFs for which the retapamulin signal is outside the start, when considering in the RPF (TIS-Ribo-Seq data) at the predicted start codon including one codon upstream and downstream of it (i.e. in total 9 nt). The red counts represent the counts from TIS-Ribo-Seq and blue from the Ribo-Seq.

**Figure S2.**
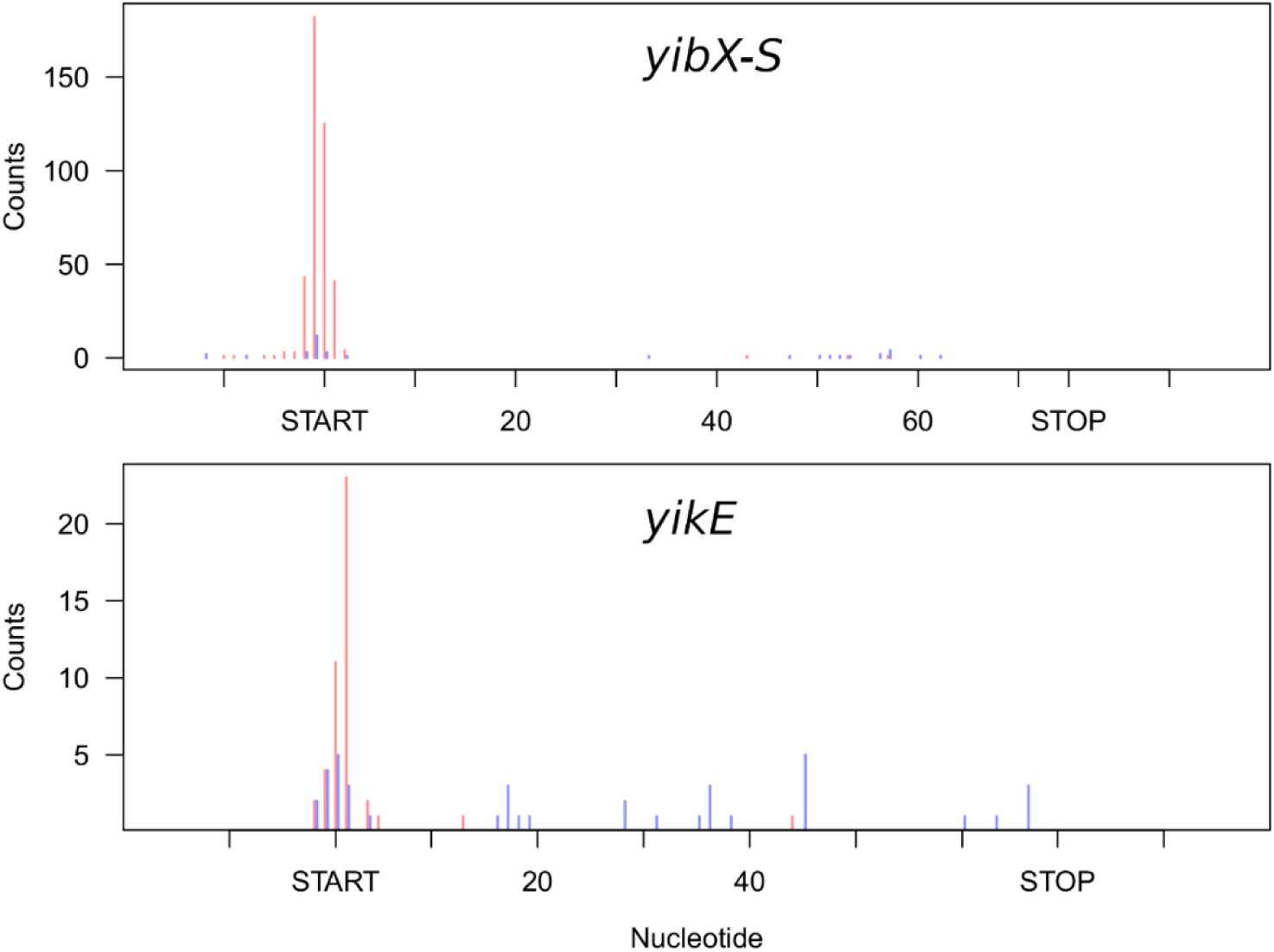
Examples of two smORFs identified using manual assessment of TISs (22) which were also successfully detected by smORFer. The red counts represent the counts from TIS-Ribo-Seq and blue from the Ribo-Seq.

## References

1. Fickett, J.W. (1982) Recognition of protein coding regions in DNA sequences. Nucl Acids Res, 10, 5303–5318.

2. Basrai, M.A., Hieter, P. and Boeke, J.D. (1997) Small open reading frames: beautiful needles in the haystack. Genome Res, 7, 768–771.

3. Maeda, N., Kasukawa, T., Oyama, R., Gough, J., Frith, M., Engstrom, P.G., Lenhard, B., Aturaliya, R.N., Batalov, S., Beisel, K.W. et al. (2006) Transcript annotation in FANTOM3: mouse gene catalog based on physical cDNAs. PLoS Genet, 2, e62.

4. Angiuoli, S.V., Gussman, A., Klimke, W., Cochrane, G., Field, D., Garrity, G., Kodira, C.D., Kyrpides, N., Madupu, R., Markowitz, V. et al. (2008) Toward an online repository of Standard Operating Procedures (SOPs) for (meta)genomic annotation. OMICS, 12, 137–141.

5. Ramamurthi, K.S. and Storz, G. (2014) The small protein floodgates are opening; now the functional analysis begins. BMC Biol, 12, 96.

6. Andrews, S.J. and Rothnagel, J.A. (2014) Emerging evidence for functional peptides encoded by short open reading frames. Nat Rev Genet, 15, 193–204.

7. Anderson, D.M., Anderson, K.M., Chang, C.L., Makarewich, C.A., Nelson, B.R., McAnally, J.R., Kasaragod, P., Shelton, J.M., Liou, J., Bassel-Duby, R. et al. (2015) A micropeptide encoded by a putative long noncoding RNA regulates muscle performance. Cell, 160, 595–606.

8. Chen, J., Brunner, A.D., Cogan, J.Z., Nunez, J.K., Fields, A.P., Adamson, B., Itzhak, D.N., Li, J.Y., Mann, M., Leonetti, M.D. et al. (2020) Pervasive functional translation of noncanonical human open reading frames. Science, 367, 1140–1146.

9. D’Lima, N.G., Ma, J., Winkler, L., Chu, Q., Loh, K.H., Corpuz, E.O., Budnik, B.A., Lykke-Andersen, J., Saghatelian, A. and Slavoff, S.A. (2017) A human microprotein that interacts with the mRNA decapping complex. Nat Chem Biol, 13, 174–180.

10. Jackson, R., Kroehling, L., Khitun, A., Bailis, W., Jarret, A., York, A.G., Khan, O.M., Brewer, J.R., Skadow, M.H., Duizer, C. et al. (2018) The translation of non-canonical open reading frames controls mucosal immunity. Nature, 564, 434–438.

11. Kondo, T., Plaza, S., Zanet, J., Benrabah, E., Valenti, P., Hashimoto, Y., Kobayashi, S., Payre, F. and Kageyama, Y. (2010) Small peptides switch the transcriptional activity of Shavenbaby during Drosophila embryogenesis. Science, 329, 336–339.

12. Matsumoto, A., Pasut, A., Matsumoto, M., Yamashita, R., Fung, J., Monteleone, E., Saghatelian, A., Nakayama, K.I., Clohessy, J.G. and Pandolfi, P.P. (2017) mTORC1 and muscle regeneration are regulated by the LINC00961-encoded SPAR polypeptide. Nature, 541, 228–232.

13. Nelson, B.R., Makarewich, C.A., Anderson, D.M., Winders, B.R., Troupes, C.D., Wu, F., Reese, A.L., McAnally, J.R., Chen, X., Kavalali, E.T. et al. (2016) A peptide encoded by a transcript annotated as long noncoding RNA enhances SERCA activity in muscle. Science, 351, 271–275.

14. Araujo-Bazan, L., Ruiz-Avila, L.B., Andreu, D., Huecas, S. and Andreu, J.M. (2016) Cytological Profile of Antibacterial FtsZ Inhibitors and Synthetic Peptide MciZ. Front Microbiol, 7, 1558.

15. Bobrovskyy, M. and Vanderpool, C.K. (2014) The small RNA SgrS: roles in metabolism and pathogenesis of enteric bacteria. Front Cell Infect Microbiol, 4, 61.

16. Ebmeier, S.E., Tan, I.S., Clapham, K.R. and Ramamurthi, K.S. (2012) Small proteins link coat and cortex assembly during sporulation in Bacillus subtilis. Mol Microbiol, 84, 682–696.

17. Hobbs, E.C., Yin, X., Paul, B.J., Astarita, J.L. and Storz, G. (2012) Conserved small protein associates with the multidrug efflux pump AcrB and differentially affects antibiotic resistance. Proc Natl Acad Sci USA, 109, 16696–16701.

18. Hobbs, E.C., Fontaine, F., Yin, X. and Storz, G. (2011) An expanding universe of small proteins. Curr Opin Microbiol, 14, 167–173.

19. Meydan, S., Marks, J., Klepacki, D., Sharma, V., Baranov, P.V., Firth, A.E., Margus, T., Kefi, A., Vazquez-Laslop, N. and Mankin, A.S. (2019) Retapamulin-Assisted Ribosome Profiling Reveals the Alternative Bacterial Proteome. Mol Cell, 74, 481–493 e486.

20. Modell, J.W., Kambara, T.K., Perchuk, B.S. and Laub, M.T. (2014) A DNA damage-induced, SOS-independent checkpoint regulates cell division in Caulobacter crescentus. PLoS Biol, 12, e1001977.

21. Salazar, M.E., Podgornaia, A.I. and Laub, M.T. (2016) The small membrane protein MgrB regulates PhoQ bifunctionality to control PhoP target gene expression dynamics. Mol Microbiol, 102, 430–445.

22. Weaver, J., Mohammad, F., Buskirk, A.R. and Storz, G. (2019) Identifying Small Proteins by Ribosome Profiling with Stalled Initiation Complexes. mBio, 10.

23. Ingolia, N.T., Ghaemmaghami, S., Newman, J.R. and Weissman, J.S. (2009) Genome-wide analysis in vivo of translation with nucleotide resolution using ribosome profiling. Science, 324, 218–223.

24. Chun, S.Y., Rodriguez, C.M., Todd, P.K. and Mills, R.E. (2016) SPECtre: a spectral coherence--based classifier of actively translated transcripts from ribosome profiling sequence data. BMC Bioinformatics, 17, 482.

25. Ingolia, N.T., Lareau, L.F. and Weissman, J.S. (2011) Ribosome profiling of mouse embryonic stem cells reveals the complexity and dynamics of mammalian proteomes. Cell, 147, 789–802.

26. Xiao, Z., Huang, R., Xing, X., Chen, Y., Deng, H. and Yang, X. (2018) De novo annotation and characterization of the translatome with ribosome profiling data. Nucl Acids Res, 46, e61.

27. Guttman, M., Russell, P., Ingolia, N.T., Weissman, J.S. and Lander, E.S. (2013) Ribosome profiling provides evidence that large noncoding RNAs do not encode proteins. Cell, 154, 240–251.

28. Aspden, J.L., Eyre-Walker, Y.C., Phillips, R.J., Amin, U., Mumtaz, M.A., Brocard, M. and Couso, J.P. (2014) Extensive translation of small Open Reading Frames revealed by Poly-Ribo-Seq. Elife, 3, e03528.

29. Fields, A.P., Rodriguez, E.H., Jovanovic, M., Stern-Ginossar, N., Haas, B.J., Mertins, P., Raychowdhury, R., Hacohen, N., Carr, S.A., Ingolia, N.T. et al. (2015) A Regression-Based Analysis of Ribosome-Profiling Data Reveals a Conserved Complexity to Mammalian Translation. Mol Cell, 60, 816–827.

30. Hsu, P.Y., Calviello, L., Wu, H.L., Li, F.W., Rothfels, C.J., Ohler, U. and Benfey, P.N. (2016) Super-resolution ribosome profiling reveals unannotated translation events in Arabidopsis. Proc Natl Acad Sci USA, 113, E7126–E7135.

31. Nakahigashi, K., Takai, Y., Kimura, M., Abe, N., Nakayashiki, T., Shiwa, Y., Yoshikawa, H., Wanner, B.L., Ishihama, Y. and Mori, H. (2016) Comprehensive identification of translation start sites by tetracycline-inhibited ribosome profiling. DNA Res, 23, 193–201.

32. Stern-Ginossar, N., Weisburd, B., Michalski, A., Le, V.T., Hein, M.Y., Huang, S.X., Ma, M., Shen, B., Qian, S.B., Hengel, H. et al. (2012) Decoding human cytomegalovirus. Science, 338, 1088–1093.

33. Bazzini, A.A., Johnstone, T.G., Christiano, R., Mackowiak, S.D., Obermayer, B., Fleming, E.S., Vejnar, C.E., Lee, M.T., Rajewsky, N., Walther, T.C. et al. (2014) Identification of small ORFs in vertebrates using ribosome footprinting and evolutionary conservation. EMBO J, 33, 981–993.

34. Calviello, L., Mukherjee, N., Wyler, E., Zauber, H., Hirsekorn, A., Selbach, M., Landthaler, M., Obermayer, B. and Ohler, U. (2016) Detecting actively translated open reading frames in ribosome profiling data. Nat Methods, 13, 165–170.

35. Shell, S.S., Wang, J., Lapierre, P., Mir, M., Chase, M.R., Pyle, M.M., Gawande, R., Ahmad, R., Sarracino, D.A., Ioerger, T.R. et al. (2015) Leaderless Transcripts and Small Proteins Are Common Features of the Mycobacterial Translational Landscape. PLoS Genet, 11, e1005641.

36. Storz, G., Wolf, Y.I. and Ramamurthi, K.S. (2014) Small proteins can no longer be ignored. Annu Rev Biochem, 83, 753–777.

37. Eastman, G., Smircich, P. and Sotelo-Silveira, J.R. (2018) Following Ribosome Footprints to Understand Translation at a Genome Wide Level. Comput Struct Biotechnol J, 16, 167–176.

38. Li, G.W., Oh, E. and Weissman, J.S. (2012) The anti-Shine-Dalgarno sequence drives translational pausing and codon choice in bacteria. Nature, 484, 538–541.

39. Del Campo, C., Bartholomaus, A., Fedyunin, I. and Ignatova, Z. (2015) Secondary Structure across the Bacterial Transcriptome Reveals Versatile Roles in mRNA Regulation and Function. PLoS Genet, 11, e1005613.

40. Langmead, B., Trapnell, C., Pop, M. and Salzberg, S.L. (2009) Ultrafast and memory-efficient alignment of short DNA sequences to the human genome. Genome Biol, 10, R25.

41. Quinlan, A.R. and Hall, I.M. (2010) BEDTools: a flexible suite of utilities for comparing genomic features. Bioinformatics, 26, 841–842.

42. Baek, J., Lee, J., Yoon, K. and Lee, H. (2017) Identification of Unannotated Small Genes in Salmonella. G3 (Bethesda), 7, 983–989.

43. Hecht, A., Glasgow, J., Jaschke, P.R., Bawazer, L.A., Munson, M.S., Cochran, J.R., Endy, D. and Salit, M. (2017) Measurements of translation initiation from all 64 codons in E. coli. Nucl Acids Res, 45, 3615–3626.

44. Bartholomaus, A. and Ignatova, Z. (2020) Codon resolution analysis of ribosome profiling data. Meth Mol Biol, in press.

45. Orr, M.W., Mao, Y., Storz, G. and Qian, S.B. (2020) Alternative ORFs and small ORFs: shedding light on the dark proteome. Nucl Acids Res, 48, 1029–1042.

46. Shabalina, S.A., Ogurtsov, A.Y. and Spiridonov, N.A. (2006) A periodic pattern of mRNA secondary structure created by the genetic code. Nucl Acids Res, 34, 2428–2437.

47. Kertesz, M., Wan, Y., Mazor, E., Rinn, J.L., Nutter, R.C., Chang, H.Y. and Segal, E. (2010) Genome-wide measurement of RNA secondary structure in yeast. Nature, 467, 103–107.

48. Ding, Y., Tang, Y., Kwok, C.K., Zhang, Y., Bevilacqua, P.C. and Assmann, S.M. (2014) In vivo genome-wide profiling of RNA secondary structure reveals novel regulatory features. Nature, 505, 696–700.

49. Wan, Y., Qu, K., Zhang, Q.C., Flynn, R.A., Manor, O., Ouyang, Z., Zhang, J., Spitale, R.C., Snyder, M.P., Segal, E. et al. (2014) Landscape and variation of RNA secondary structure across the human transcriptome. Nature, 505, 706–709.

50. Tiwari, S., Ramachandran, S., Bhattacharya, A., Bhattacharya, S. and Ramaswamy, R. (1997) Prediction of probable genes by Fourier analysis of genomic sequences. Comput Appl Biosci, 13, 263–270.

51. Woolstenhulme, C.J., Guydosh, N.R., Green, R. and Buskirk, A.R. (2015) High-precision analysis of translational pausing by ribosome profiling in bacteria lacking EFP. Cell Rep, 11, 13–21.

